# Automatic and accurate ligand structure determination guided by cryo-electron microscopy maps

**DOI:** 10.1101/2022.08.16.504149

**Authors:** Andrew Muenks, Samantha Zepeda, Guangfeng Zhou, David Veesler, Frank DiMaio

## Abstract

Advances in cryo-electron microscopy (cryoEM) and deep-learning guided protein structure prediction have expedited structural studies of protein complexes. However, methods for accurately determining ligand conformations are lacking. In this manuscript, we develop a tool for automatically determining ligand structures guided by medium-resolution cryoEM density. We show this method is robust at predicting ligands in maps as low as 6Å resolution, and is able to correct receptor sidechain errors. Combining this with a measure of placement confidence, and running on all protein/ligand structures in EMDB, we show that 58% of ligands replicate the deposited model, 16% confidently find alternate conformations, 22% have ambiguous density where multiple conformations might be present, and 4% are incorrectly placed. For five cases where our approach finds an alternate conformation with high confidence, high-resolution crystal structures validate our placement. This tool and the resulting analysis should prove critical in using cryoEM to investigate protein-ligand complexes.

## INTRODUCTION

Recent advancements in both microscope hardware and computational processing have led to cryo-electron microscopy (cryoEM) emerging as a mainstream method for biomolecular structure determination. While in ideal cases cryoEM data approaches atomic resolutions [1, 2, 3], most structures determined by cryoEM are in the 3–5Å resolution range. At these resolutions, model building is time consuming, error prone, and often ambiguous. To assist this process, methods have been developed to automatically build de novo polypeptide chains into EM data [4–7], and with the advent of AlphaFold 2, high-quality starting models can oftentimes be obtained from sequence information alone [8, 9]. While these methods help build protein models into cryoEM density, tools for automatic fitting of small molecule ligands into cryoEM data are limited. Given the widespread adoption of cryoEM in academia and in industry to support translational studies of drug targets, the ability to accurately model ligand-bound structures is paramount.

There are numerous automated tools from X-ray crystallography for modeling small molecule ligands [10–13]. However, their methodology often precludes their use for interpreting all but the highest-resolution cryoEM maps. Traditional ligand fitting methods rely on shape and topological features of density maps to match [10] or build [11, 12, 13] the ligand into density. But as resolution decreases below 3Å, the topological features these methods rely on become less defined, and their accuracy in modeling ligands within 1Å RMSD of a reference ligand falls below 20% [13, 14]. Along with map features, chemical force fields have provided an energetic approach to accurately fit ligands into their respective density. Two approaches — GemSpot [15] and MDFF [16] — build upon the ligand-docking software GLIDE to model ligands into cryoEM data. However, both require user input in either selecting models or choosing a starting configuration, limiting the automation and applicability of these approaches.

The protein modeling software Rosetta recently incorporated a new small molecule force field, RosettaGenFF, which accurately models the energetics of arbitrary biomolecules in a manner balanced against Rosetta’s protein forcefield [17]. Combining this energy model with a genetic-algorithm (GA) optimization method allowing for full receptor sidechain flexibility, GALigandDock, yielded superior performance in ligand docking accuracy compared to other state-of-the-art methods. Here, we leverage the docking power of RosettaGenFF and GA optimization to overcome the challenges of modeling small molecules at near-atomic resolution. We integrated cryoEM density data with the physically realistic force field of RosettaGenFF to create RosettaEMERALD (EM Maps ERoded for Automatic Ligand Docking) for robust ligand modeling into cryoEM maps. We evaluated the performance of EMERALD on all non-ion-mediated ligand-bound protein structures deposited in the EMDB [18] (see *Methods*) and compared our results to their respective deposited structures and high-resolution crystal structures when available.

## RESULTS

An overview of EMERALD is illustrated in Figure 1. GALigandDock places ligands in a protein pocket by iteratively refining a pool of 100 conformations, selecting the best 100 models at each generation using predicted energy. To enable this method to use cryoEM density, two changes were integral: density-guided initial ligand placement and the use of density in model selection at each round. Our initial placement (fully described in *Methods*) first models density as a pseudo-atomic skeleton (Fig. 1b). When generating the initial population of ligands, ligands are placed at the center of the skeleton and restrained to points in the skeleton. At each iteration, the population of ligand conformers along with their surrounding flexible side chains are further optimized against the sum of a weighted density correlation and the RosettaGenFF energy (Fig. 1c) and finally refined in Rosetta to minimize the energy of the models (Fig. 1d). The full protocol generates a structure in 30–120 minutes, depending on the size of the ligand and the cryoEM map.

**Figure 1.**
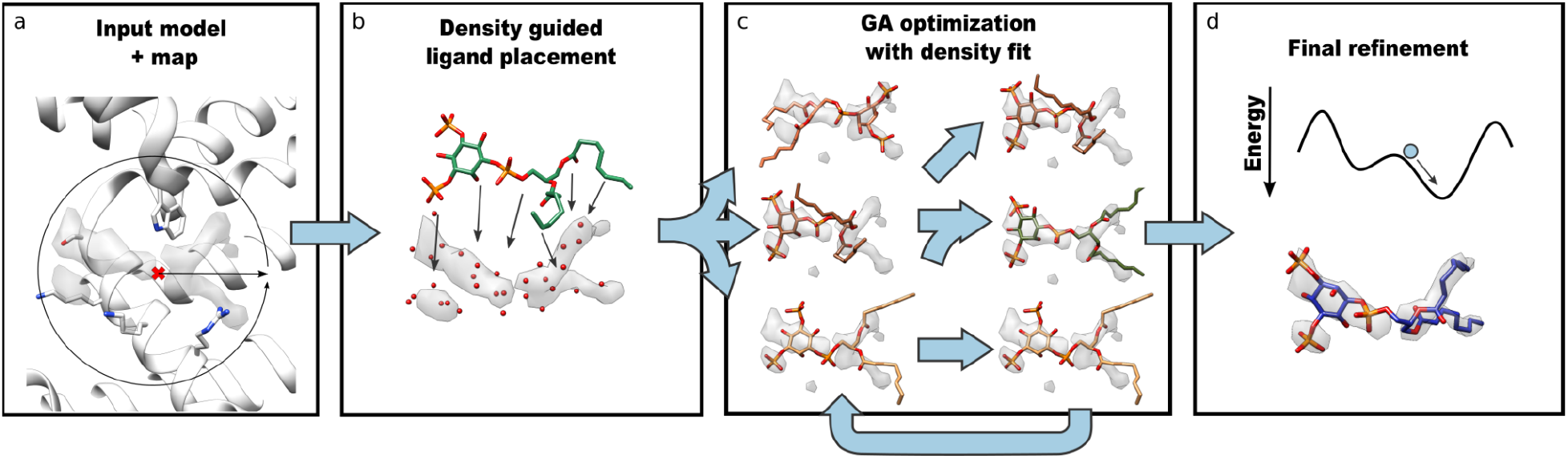
Overview of EMERALD docking protocol. (a) The cryoEM map, coordinates of the receptor, and the location of the binding site are provided as inputs. (b) All unmodeled density in the pocket is converted to a pseudo-atomic skeleton (independent of ligand identity), which is used to generate an initial set of ligand conformers. (c) Using a genetic algorithm, the pool of ligand conformers are optimized against Rosetta energy and density fit. (d) The 20 poses with lowest energy are refined in Rosetta.

To test RosettaEMERALD, we ran our docking protocol on all ligands with 25 or fewer rotatable torsion angles present in deposited cryoEM structures determined at a minimum of 6Å resolution. This yielded 1053 ligands to be placed. For each model, we ran three independent trajectories, and we analyzed the resulting models using three different criteria: a) agreement of the deposited model to the lowest energy predicted structure; b) density fit and number of protein/ligand hydrogen bonds; and c) convergence of the three trajectories. This last criterion is used to evaluate the confidence in a predicted model.

The results of these docking trajectories are summarized in Figure 2. In 58% of the cases, our density-guided docking produced a top model within 1Å RMSD (considering all non-hydrogen atoms in the ligand) of the deposited model after energy minimization (“match”, Fig 2a.). While an RMSD cutoff of 2Å has traditionally been used for docking success, the lack of confidence in the low resolution reference models and inability of RMSD to consider receptor contacts led us to divide results further by *density correlation* and *hydrogen bond contacts*. There were 398 cases (38%) where EMERALD produced a model with an RMSD value greater than 1Å, and the model was similar or better than the deposited model in both metrics (“non-match, similar or better quality”). The smallest group belonged to 47 cases where the deposited model was not recapitulated, but the EMERALD model had a worse density fit or fewer hydrogen bonds than the deposited model (“non-match, worse quality”, 4%). We found that incorporating EM data in GALigandDock is necessary for recapitulating deposited ligand structures with high success rates (Suppl. Fig. 1). Also, docking success is resilient to changes in overall map resolution (Suppl. Fig. 2a-c) and docking performs worse as the flexibility of the ligand increases (Suppl. Fig. 2d-f).

**Figure 2.**
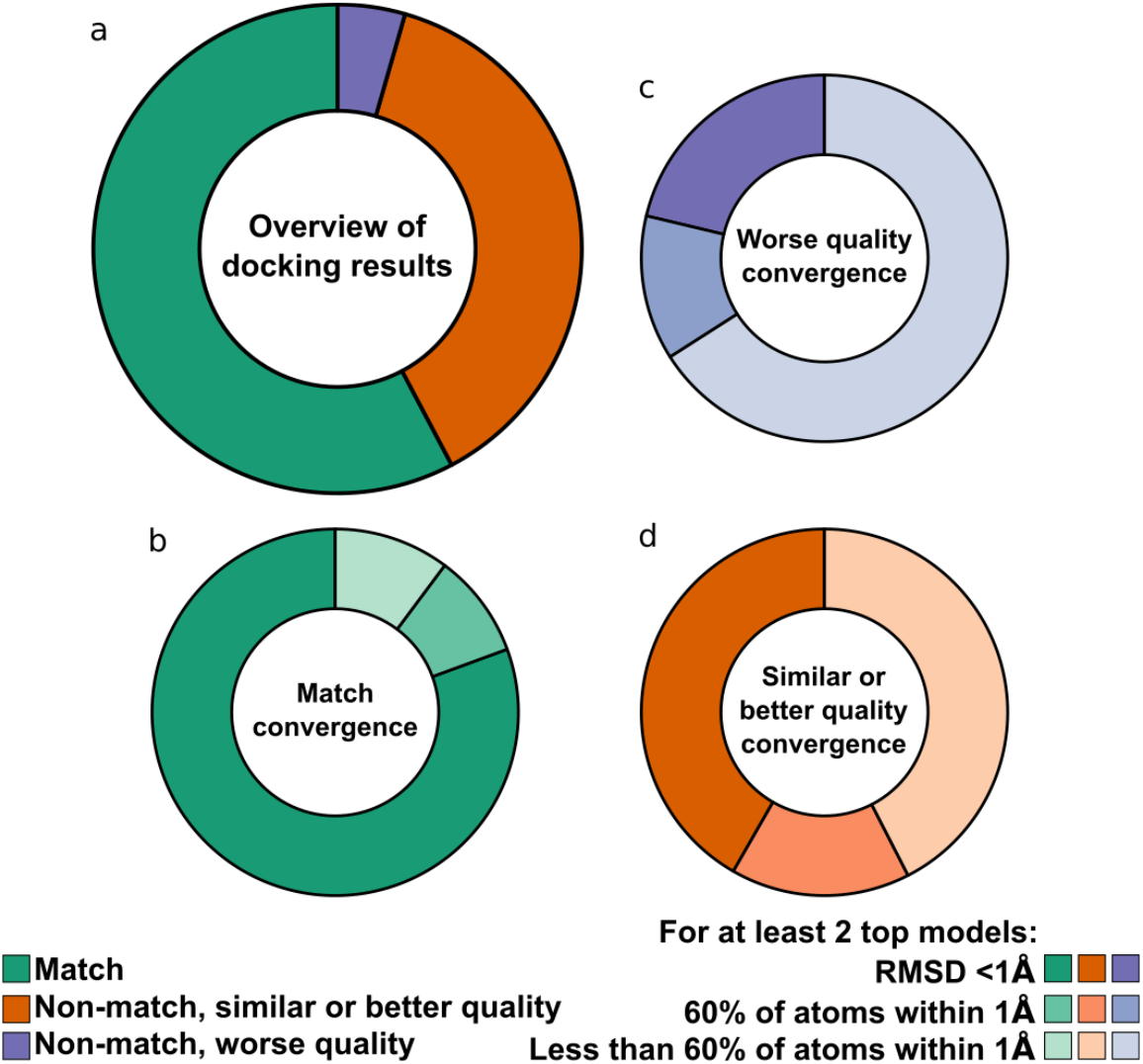
Benchmarking EMERALD against the EMDB. (a) A comparison of EMERALD models to the deposited structures over 1053 EMDB-deposited complexes. In total, 58% of EMERALD-docked models were placed within 1Å RMSD of the deposited ligand (“match”, *green*); 38% were more than 1Å RMSD of the deposited ligand but had similar or better density correlations and numbers of hydrogen bonds (“similar or better quality”, *orange*), and 4% were 1Å RMSD of the deposited ligand and had worse density correlations or number of hydrogen bonds (“worse quality”, *blue*) (b-d) The convergence of the best ranking models across multiple runs for matches (b), worse quality (c), and similar or better quality (d) cases. The darkest shade had multiple runs converge with all atoms within 1Å of each other, the middle shade had multiple runs converge with at least 60% of atoms within 1Å, and the lightest shade had divergent top-scoring models.

Because of the low-resolution of the density maps, it is difficult to interpret the quality of docked poses from density fit and receptor interactions alone. To instill more confidence in docking results, we analyzed the convergence among the top ranked ligand poses across three replicates (Fig. 2b-d). Of the cases within 1Å RMSD, 2 or more of the trajectories converge for 81% of cases, further strengthening the quality of the matched cases (Fig. 2b). Moreover, only 21% of the worse-quality cases converge on the same ligand model (Fig. 2c). Given how well trajectory convergence agrees with these categories, it can serve as a proxy for confidence when our docked model differs from the reference model in ambiguous cases. 42% of the ambiguous cases have similar top models across our trajectories (Fig. 2d), giving us confidence in an alternative model to the deposited structure for those entries.

Our dataset includes 15 of the 20 cases benchmarked for the GemSpot pipeline, with 5 cases filtered out of the dataset for being peptides or having inter-residue bonds like ion coordination. For 13 of the 15 ligands, EMERALD produced a ligand within 1Å of the deposited structure, with 9 of those placements assessed as confident. For the other two cases, our models disagreed with the deposited model, and GemSpot also found solutions different from the deposited model in these two cases [15].

### Crystal models confirm alternate conformations for EM data

To cross-validate our results — particularly in cases where we found a different solution than the deposited model — we looked for all models with a corresponding high-resolution crystal structure (see *Methods*). We identified 100 cases where EMERALD converged on a ligand placement and a corresponding high-resolution crystal structure was available. The converged docked model was within 1Å RMSD of the ligand modeled in the crystal structure for 67% of cases, while 58% of the deposited EM models were within this distance. Considering cases where the model predicted from EMERALD and the reference EM model differ, there were 6 cases where the EMERALD model was within 1.0Å RMSD to the crystal structure while the EM model was not, 3 cases where the EM model was within 1.0Å of the crystal structure but the EMERALD model was not, and 8 cases where both models differed from the crystal structure by more than 1.0Å. Additionally in 5 of the 6 cases where our model predicts the crystal structure, our ligand model improves density correlation by at least 0.03, compared to the deposited cryoEM model.

We show docked models supported by crystal structures in Figure 3 to highlight the quality of our protocol. These examples include: a) the hippocampal AMPA receptor with the antagonist MPQX (EMDB: 23292, PDB: 7LEP) [19], where are model makes additional hydrogen bond and π-stacking interactions with the ligand, matching the crystal structure [20]; b) NBQX in an AMPA receptor (EMDB: 12805, PDB: 7OCE)[21], where the ligand is flipped, better matching the density, and making bidentate interactions with a nearby arginine residue (Fig. 3b); c) DNMDP bound to the SLFN12-PDE3A complex (EMDB: 23495, PDB: 7LRD)[22], where small changes better match the crystal structure (Figure 3c); d) an ADP molecule in ClpB disaggregase (EMDB: 21553, PDB: 6W6E)[23] (Fig. 3d), where the phosphate groups recapitulate the crystal structure (Fig. 3d); and e) a glutamate ligand in the AMPA glutamate receptor (EMDB: 12806, PDB: 7OCF) [21], which was missing an oxygen atom in the deposited structure; when the full glutamate molecule is docked, the carboxylates are placed in a configuration matching the crystal structure [24] (Fig. 3e).

**Fig. 3.**
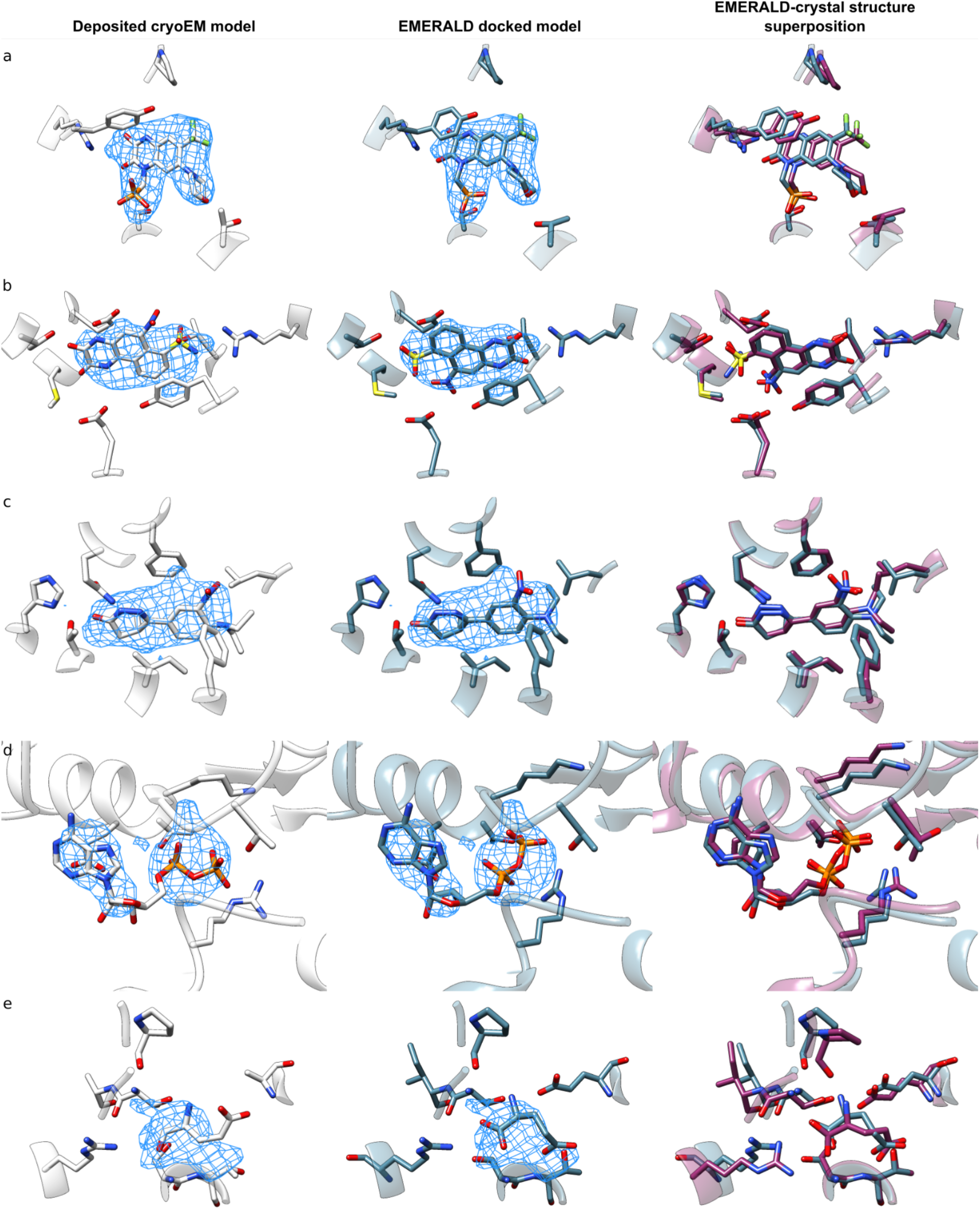
Alternate conformations predicted with EMERALD match high-resolution crystal structures. (a) Alternate conformation of antagonist ZK200775 in AMPA receptor (EMDB: 23292, PDB: 7LEP). The deposited model (left, *white*) has a phosphate group popped out of the density and tyrosine and threonine residues are not interacting with the ligand. The docked model from EMERALD (middle, *blue*) has the phosphate group fit into density and the tyrosine and threonine make the same interactions as the crystal structure (right, *purple*, PDB: 5ZG2). (b) The deposited model of NBQX in the AMPA receptor (EMDB: 12805, PDB: 7OCE) does not fully explain the observed density (left, *white*). The docked model (middle, *blue*) better matches the density, making interactions with an arginine we also see in the crystal model (right, *purple*, PDB: 6FQH). (c) The deposited model for DNMDP bound to the SLFN12-PDE3A complex (EMDB: 23495, PDB: 7LRD) poorly matches the observed density (left, *white*). The docked model (middle, *blue*) fills the unexplained density (left, *white*), superimposing closely to the crystal model (right,*purple*, PDB: 7KWE). (d) The deposited model of ADP bound to ClpB (EMDB: 21553, PDB: 6W6E) has the phosphates oriented away from a binding motif loop (left, *white*). Our docked model (middle, *blue*) adopts a different conformation placing the phosphates towards a nucleotide binding domain; the crystal structure adopts this alternate conformation (right,*purple*, PDB: 5LJ8). (e) The deposited model of an alternate conformation of glutamate in AMPA receptor (EMDB: 12806, PDB: 7OCF) is missing an oxygen atom and the amino group is not interacting with the receptor (left, *white*). A corrected docked model from EMERALD (middle, *blue*) finds a conformation nearly identical to a high-resolution crystal model of the same receptor (right, *purple*, PDB: 3TKD).

There were three cases where our docking protocol found a ligand different than the crystal structure, while the EM model matched the crystal structure closely. All three cases were different maps of the same system, a folate molecule bound to MERS-CoV (EMDBs: 20543, 20544, 23674; PDBs: 6Q05, 6Q06, 7M5E)[25,26]. In all 3, the EMERALD model and the crystal structure only differ in the placement of a flexible arm with high B-factors in the crystallographic data (Suppl. Fig. 3) (PDB: 5VYH)[27]. These results lend more support for EMERALD convergence as a confidence metric, which we used to further find instances of alternate ligand conformations.

### Docked poses reveal plausible alternate conformations

Even without crystal structures for reference, trajectory convergence and improved ligand density fit provide confidence in other docked poses. In the case of an antimicrobial bound multiple transferable resistance (Mtr) pump (EMDB: 21228, PDB: 6VKS) [28], our protocol converges on an ampicillin molecule that is flipped so that its phenyl group is now in a pocket of unassigned density (Fig. 4a, b). While the deposited model places the phenyl group sandwiched between two phenylalanine residues (Fig. 4a), our docked model packs the group near a cluster of hydrophobic residues known to interact with other antibiotics [28] (Fig. 4b). Additionally, nearby arginine, serine, and threonine residues have been suggested to generally coordinate ligands binding to the pump [28]; our model has the carboxyl group positioned to make interactions with these residues directly or possibly through bridging water molecules. While it is likely that an antibiotic would bind non-specifically to this site, EMERALD ranks our presented orientation the highest across all three trajectories, and there is a large predicted energy gap (about 10 kcal/mol) between the converged conformation and the best-scoring conformation with the phenyl group outside this hydrophobic pocket, suggesting that this pose is strongly favored by EMERALD.

**Figure 4.**
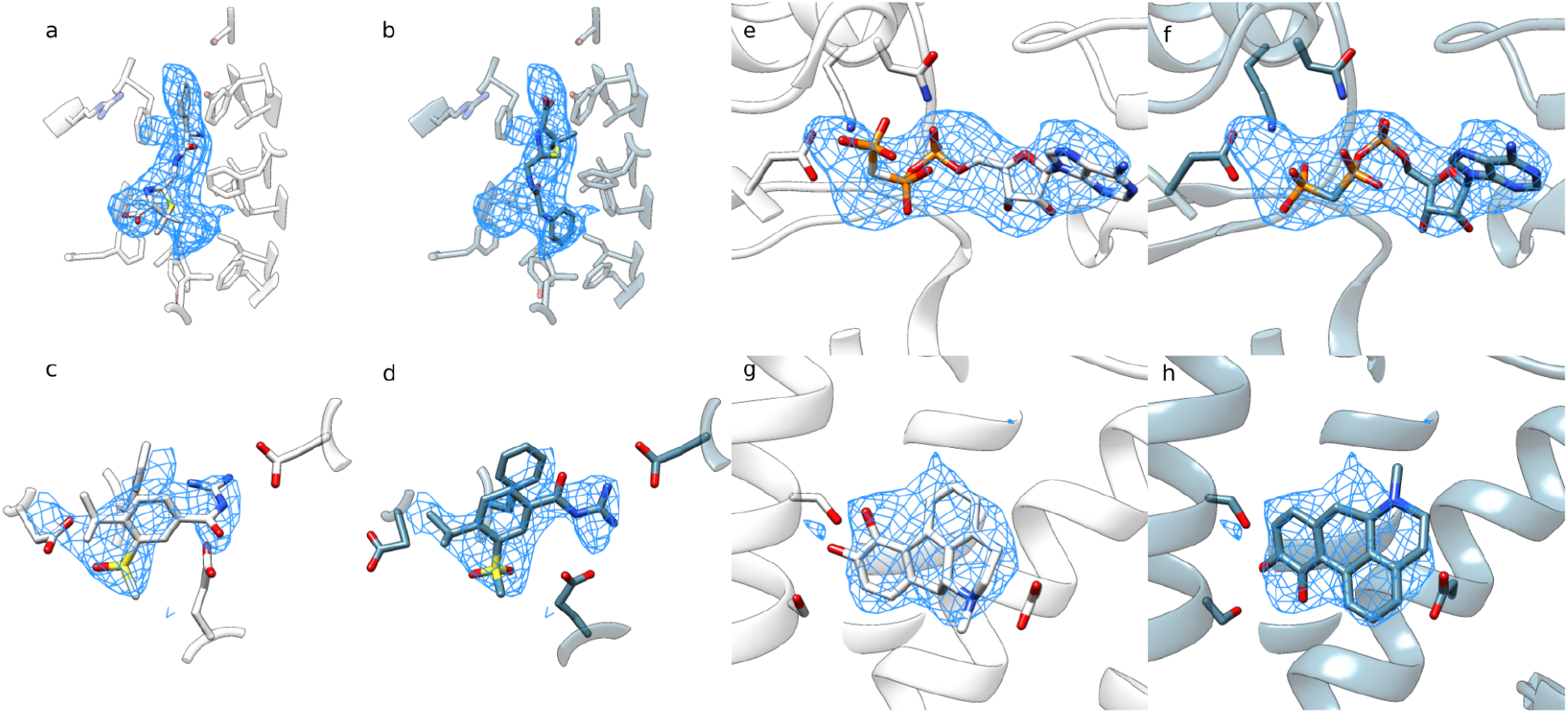
Alternate conformations found by EMERALD in cases without crystal structures. (a) The deposited structure of Ampicillin bound to Mtr pump (EMDB: 21228, PDB: 6VKS) has unexplained density in a hydrophobic pocket that is filled by the phenyl group in the docked model. (b) The EMERALD-docked model fills this missing density, making similar hydrophobic interactions to the receptor. (c) The deposited model of the binding pocket of cariporide in the Na+/H+ exchanger I (EMDB: 30849, PDB: 7DSX) has a guanidine group making cation-π interactions with a phenylalanine, but weakly interacting with a nearby aspartate and glutamate. (d) In the EMERALD-docked model, the guanidine group makes a strong salt bridge interaction with the aspartate and glutamate. (e) The deposited model of an ATP analog in flippase ATP11C (EMDB: 30163, PDB: 7BSP) has the gamma phosphate poking out of the density; (f) the EMERALD-docked model better matches the density, with the gamma phosphate anchored by a neighboring lysine. (g) The deposited model of agonist apomorphine in a dopamine receptor (EMDB: 22510, PDB: 7JVQ) explains the density well and makes largely hydrophobic interactions with the receptor. (h) The EMERALD-docked model strongly prefers an orientation where the molecule is rotated by 180 degrees.

Despite some cases not converging on lowest energy models within 1Å RMSD of each other, an important substructure of the ligand may align. One slight change in ligand and protein model comes from the Na+/H+ exchanger I inhibited by cariporide (EMDB: 30849, PDB: 7DSX) [29] (Fig. 4c, d). The original structure has the guanidine group making cationic interactions with neighboring phenylalanine, glutamate, and aspartate residues, all shown to be important for drug binding [30–33] (Fig. 4c). The docked model deviates by adopting an orientation for the guanidine group in-plane with the ligand, making more direct interactions with the aspartate and glutamate but losing its cation-π interaction with the phenylalanine residue (Fig. 4d). However, the phenylalanine now forms offset π-stacking with the ligand. The planar guanidine orientation appears in the top model across each triplicate run and over half of the non-hydrogen atoms among top models are within 1Å of each other. Furthermore, a guanidine conformation like the deposited model does not appear in the 20 lowest-energy models for any trajectory. All this suggests that this conformation is favored to one forming cation-π interactions and could explain the experimental data in a new light.

Additional cases with confident alternative models are shown in Fig. 4e-h. For the ATP analog in a structure of the ATP11C flippase (EMDB: 30163, PDB: 7BSP) [34] the gamma phosphate sticks out of density in the deposited model (Fig. 4e) but is modeled into the density and interacting with a nearby lysine residue in the docked model (Fig. 4f). Meanwhile the docked model for an apomorphine molecule in a dopamine receptor (EMDB: 22510, PDB: 7JVQ) [35] is flipped 180° and does not make additional interactions with the receptor when compared to the deposited model (Fig. 4g-h), but Rosetta converges on the same model for all 3 replicates, suggesting some level of confidence.

### Low-confidence unmatched cases show pseudo-symmetry or weak density

While our analysis confidently discovers alternate ligand models, 58% of docked molecules with similar quality to the deposited model have medium or low confidence. We found that small molecules that have pseudo-symmetry or have flexible moieties represent these low-confidence cases because of the challenges they provide from their often noisy and inconclusive density. In some instances, two or more replicates of EMERALD agree on a substructure of the molecule (EMDB: 21844, PDB: 6WLW, dark blue, Fig. 5a, b)[36], but differ in a rotamer of a functional group or a flexible group (light blue, Fig. 5a, b). For other ligands, ambiguous density leads to little agreement among the reference model and low-energy Rosetta models (Fig. 5c, d). The authors for the allosteric modulator of a dopamine receptor (EMDB: 30395, PDB: 7CKZ) note the lack of confidence in the deposited structure [37], but have mutagenesis studies to confirm the conformation modeled (Fig. 5c)[38]. However, one model found with EMERALD aligns with their opposing model and fulfills an unexplained region of density in the deposited model (Fig. 5d). Altogether, these entries show the difficulty in interpreting cryoEM data at medium to low resolution leading to ambiguous density explanations for a single map, and the limits to automated ligand docking using our protocol.

**Figure 5.**
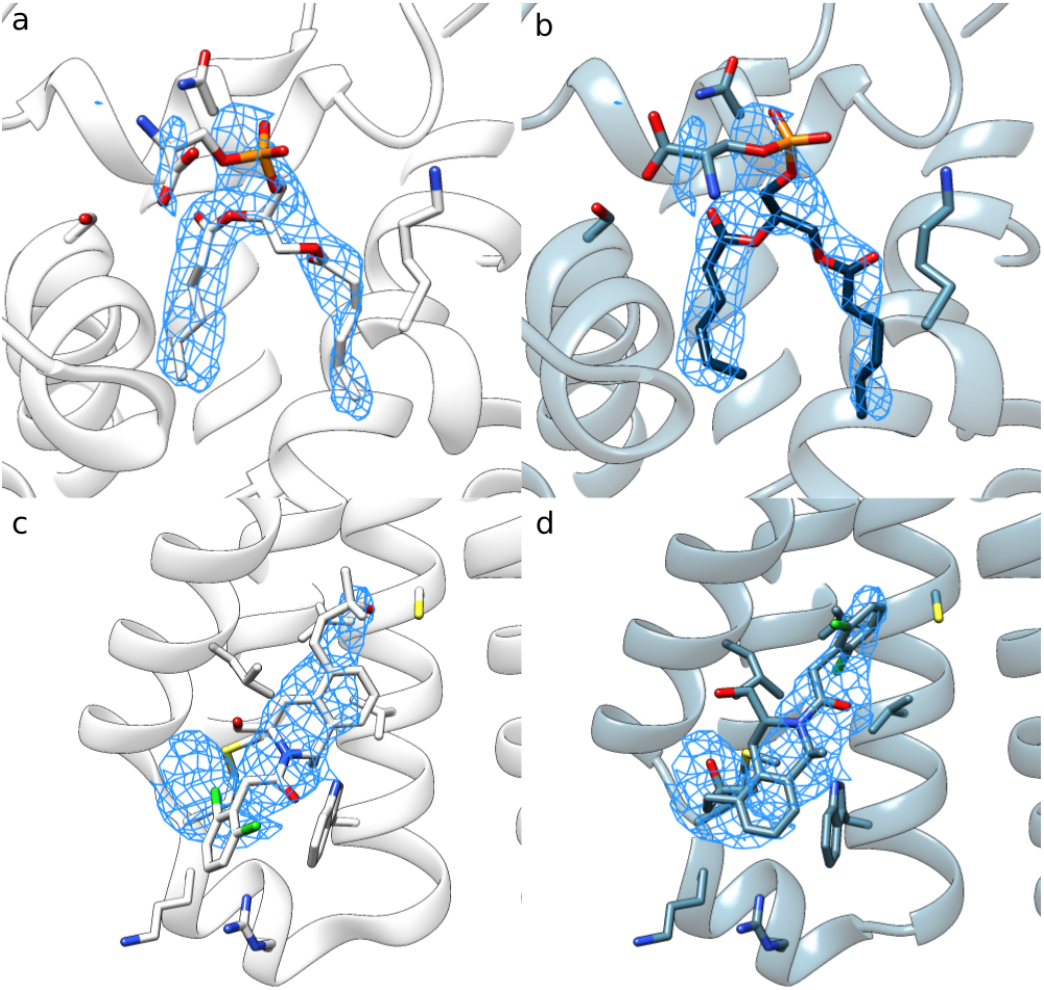
Examples of low-confidence docked models where there may be ambiguity or heterogeneity in the data. (a) The deposited model of a phosphoserine lipid in ATPase (EMDB: 21844, PDB: 6WLW) and the EMERALD-docked model (b) place the fatty acid tails in strong density, but have differences in the likely mobile head group. The lowest-energy models for all 3 triplicates find the same lipid tail orientations (*dark blue*), but differ in the head group. (c) The deposited model of LY3154207 bound to DRD1 (EMDB: 30395, PDB: 7CKZ) and EMERALD-docked model (d) adopt different conformations. With low resolution density and few residue binding partners, both models are equally plausible.

### Cases with worse ligand models show poor initial sampling

To learn what improvements could be made to EMERALD in the future, we looked at instances where EMERALD predicts a ligand with worse metrics than the reference model. We found that these cases often had density that is discontinuous or noisy, leading to incorrect skeletonization. For a ubiquinone binding electron transport protein (EMDB: 30475, PDB: 7CUW) [39], the density skeleton only finds density near the head group (Suppl. Fig. 4c). Without a complete skeleton, the initial population struggles to find the deposited conformation, placing the head group exposed to solvent (Suppl. Fig. 4b). In this case, if the 2.63Å data is instead truncated at 4.0Å resolution, the density becomes more continuous, and the skeleton generated by EMERALD matches the ligand conformation much more closely (Suppl. Fig. 4d). With a complete skeleton, the docked model is no longer worse than the deposited model. The head group of the lowest-energy model makes the same hydrogen bond interactions as the deposited model, and the docked model improves density correlation by 0.03 (Suppl. Fig. 4e). This underscores the importance of the initial sampling step, especially when evaluating ligands with a large number of rotatable bonds, and identifies areas for future upgrades in EMERALD.

### Blind modeling of linoleic acid

To demonstrate our protocol’s utility in structure determination, we used EMERALD to create a model for linoleic acid bound in a previously undetermined protein structure. Determining this model manually would be an arduous task considering high flexibility of the ligand (Fig. 6a). Despite the difficulty of modeling the suspected ligand, EMERALD predicts a small molecule conformation that fits the density, makes an anchoring electrostatic interaction with a neighboring arginine residue, and introduces little torsional strain throughout the hydrophobic tail (Fig. 6b). This placement is supported by the structure of linoleic acid bound to a related protein (EMDB: 11145, PDB: 6ZB5)[40]. Creating the model required no user input once ligand restraint files were made, and the ease and accuracy when modeling linoleic acid prove the value of EMERALD for structure determination.

**Fig. 6.**
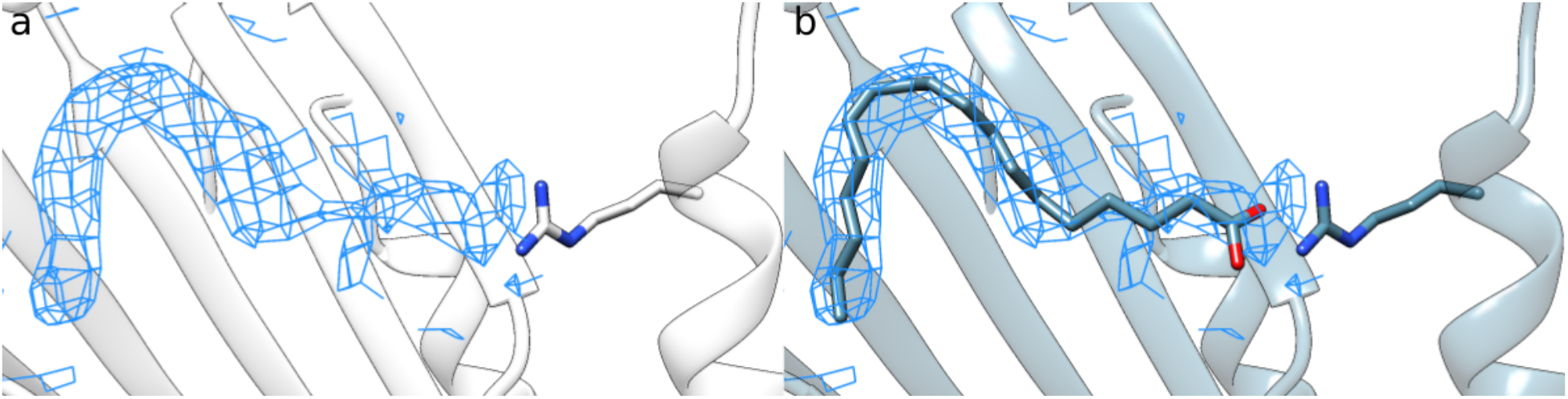
Blind modeling of linoleic acid. (a) Unmodeled density for linoleic acid. The density is difficult to fit with a long, flexible hydrophobic lipid tail of the ligand. (b) Output model from density-guided docking. The model makes an anchoring electrostatic interaction with a nearby arginine residue and models the tail strain-free into the density.

## DISCUSSION

Here, we show a method EMERALD that accurately and automatically models small molecules into cryoEM maps after being benchmarked on over one thousand ligand-bound entries in the EMDB. EMERALD identifies a confident solution in 62% of these cases, in some cases correcting errors present in the deposited model. Moreover, we show this fully automated protocol determining the conformation of linoleic acid in a previously unsolved structure.

The method should be generally applicable to most ligands with fewer than 25 rotatable bonds; larger ligands have too large of a search space for this algorithm to effectively sample. Discontinuous or noisy density also proved challenging, though modified map processing to improve density connectivity was shown to rescue at least one of these cases. Our current approach only models a single ligand at a time, which complicates density assignment for structures with ligands close together like electron transport proteins. Finally, Rosetta’s poor handling of metal ions precludes modeling ions as cofactors or as ligands themselves, leaving a significant group of proteins unanalyzed [41]

Currently, our method requires the modeler to know the identification and approximate binding location of the ligand, a non-trivial task when studying novel protein-ligand complexes. For more utility during model building, our method could expand to recognize potential unmodeled ligand blobs and quickly assess possible ligands to determine identity. As is, however, EMERALD offers an automatic tool for ligand modeling that will prove helpful for the now common scenario of ligand-bound structure determination through cryoEM, and EMERALD will serve as a valuable addition to the toolkit of Rosetta EM modeling methods [5, 42, 43] for model building under one software package.

## AVAILABILITY

All methods described are available as part of Rosetta, using weekly releases after week X, 2022. The Rosetta XML files and flags for running all the refinements discussed in this manuscript are included in Supplemental Methods.

## ACKNOWLEDGEMENTS

This study was supported by the National Science Foundation Graduate Research Fellowship under Grant No. DGE-1762114 (to A.M.), the National Institute of General Medical Sciences (1R01GM123089-01, to F.D.), the Defense Threat Reduction Agency (GRANT13030960, to F.D. and G.Z.), the National Institute of Allergy and Infectious Diseases (DP1AI158186 and HHSN272201700059C to D.V.), a Pew Biomedical Scholars Award (D.V.), an Investigators in the Pathogenesis of Infectious Disease Awards from the Burroughs Wellcome Fund (D.V.), Fast Grants (D.V.), the Bill & Melinda Gates Foundation (OPP1156262 to D.V.), the University of Washington Arnold and Mabel Beckman cryoEM center and the National Institute of Health grant S10OD032290 (to D.V.), the National Institute of General Medical Sciences (5T32GM008268-32 to S.K.Z.). D.V. is an Investigator of the Howard Hughes Medical Institute. Any opinion, findings, and conclusions or recommendations expressed in this material are those of the authors(s) and do not necessarily reflect the views of the National Science Foundation.

## REFERENCES

1. Yip, K.M., Fischer, N., Paknia, E., Chari, A. & Holger, S. Atomic-resolution protein structure determination by cryo-EM. Nature 587, 157–161 (2020).

2. Nakane, T. et al. Single-particle cryo-EM at atomic resolution. Nature 587, 152–156 (2020).

3. Merk, A. et al. 1.8 Å resolution structure of β-galactosidase with a 200 kV CRYO ARM electron microscope. IUCrJ 7, 639–643 (2020).

4. Wang, R.Y.R. et al. De novo protein structure determination from near-atomic-resolution cryo-EM maps. Nat Methods 12, 335–338 (2015).

5. Terwilliger, T.C., Adams, P.D., Afonine, P.V. & Sobolev, O.V. Cryo-EM map interpretation and protein model-building using iterative map segmentation. Protein Science 29, 87–99 (2020).

6. Terashi, G. & Kihara, D. De novo main-chain modeling for EM maps using MAINMAST. Nat Commun 9, 1618 (2018).

7. He, J. & Huang, S.Y. Full-length de novo protein structure determination from cryo-EM maps using deep learning. Bioinformatics 37, 3480–3490 (2021).

8. Jumper, J. et al. Highly accurate protein structure prediction with AlphaFold. Nature 596, 583–589 (2021).

9. Baek, M. et al. Accurate prediction of protein structures and interactions using a three-track neural network. Science 373, 871–876 (2021).

10. Emsley, P. & Cowtan, K. Coot: model-building tools for molecular graphics. Acta Cryst. D60, 2126–2132 (2004).

11. Oldfield, T.J. X-LIGAND: an application for the automated addition of flexible ligands into electron density. Acta Cryst. D57, 696–705 (2001).

12. Zwart, P.H., Langer, G.G. & Lamzin, V.S. Modelling bound ligands in protein crystal structures. Acta Cryst. D60, 2230–2239 (2004).

13. Terwilliger, T.C., Klei, H., Adams, P.D., Moriarty, N.W. & Cohn, J.D. Automated ligand fitting by core-fragment fitting and extension into density. Acta Cryst. D62, 915–922 (2006).

14. Evrard, G.X., Langer, G.G., Perrakis, A. & Lamzin, V.S. Assessment of automatic ligand building in ARP/wARP. Acta Cryst. D63, 108–117 (2007).

15. Robertson, M.J., van Zundert, G.C.P., Borrelli, K. & Skiniotis, G. GemSpot: A pipeline for robust modeling of ligands into cryo-EM maps. Structure 28, 707–716 (2020).

16. Vant, J.W. et al. Flexible fitting of small-molecules into electron microscopy maps using molecular dynamics simulations with neural network potentials. J. Chem. Inf. Model 60, 2591–2604 (2020).

17. Park, H., Zhou, G., Baek, M., Baker, D. & DiMaio, F. Force field optimization guided by small molecule crystal lattice data enables consistent sub-angstrom protein–ligand docking. J. Chem. Theory Comput. 17, 2000–2010 (2021).

18. Lawson, C.L., et al. EMDataBank unified data resource for 3DEM. Nucleic Acids Res. 44, D396–D403 (2016).

19. Yu, J. et al. Hippocampal AMPA receptor assemblies and mechanism of allosteric inhibition. Nature 594, 448–453 (2021).

20. Sobolevsky, A.I., Rosconi, M.P. & Gouaux, E. X-ray structure of AMPA-subtype glutamate receptor: symmetry and mechanism. Nature 462, 745–756 (2009).

21. Zhang, D., Watson, J.F., Matthews, P.M., Cais, O. & Greger, I.H. Gating and modulation of a hetero-octameric AMPA glutamate receptor. Nature 594, 454–458 (2021).

22. Garvie, C.W. et al. Structure of PDE3A-SLFN12 complex reveals requirements for activation of SLFN12 RNase. Nat. Commun. 12, 4375 (2021).

23. Yin, Y. et al. Structural basis for aggregate dissolution and refolding by the Mycobacterium tuberculosis ClpB-DnaK bi-chaperone system. Cell Rep. 35, 109166 (2021).

24. Krintel, C. et al. Thermodynamics and structural analysis of positive allosteric modulation of the ionotropic glutamate receptor GluA2. Biochem J. 441, 173–178 (2012).

25. Park, Y.J. et al. Structures of MERS-CoV spike glycoprotein in complex with sialoside attachment receptors. Nat. Struct. Mol. Biol. 26, 1151–1157 (2019).

26. Sauer, M.M. et al. Structural basis for broad coronavirus neutralization. Nat. Struct. Mol. Biol. 28, 478–486 (2021).

27. Pallesen, J. et al. Immunogenicity and structures of a rationally designed prefusion MERS-CoV spike antigen. Proc. Natl. Acad. Sci. USA 114, E7348–E7357 (2017).

28. Lyu, M. et al. Cryo-EM structures of a gonococcal multidrug efflux pump illuminate a mechanism of drug recognition and resistance. mBio 11, e00996–20 (2020).

29. Dong, Y. et al. Structure and mechanism of the human NHE1-CHP1 complex. Nat. Commun. 12, 3474 (2021).

30. Touret, N., Poujeol, P. & Counillon, L. Second-site revertants of a low-sodium-affinity mutant of the Na+/H+ exchanger reveal the participation of TM4 into a highly constrained sodium-binding site. Biochemistry 40, 5095–5101 (2001).

31. Noël, J., Germain, D. & Vadnais, J. Glutamate 346 of human Na+-H+ exchanger NHE1 is crucial for modulating both the affinity for Na+ and the interaction with amiloride derivatives. Biochemistry 42, 15361–15368 (2003).

32. Khadilkar, A., Iannuzzi, P. & Orlowski, J. Identification of sites in the second exomembrane loop and ninth transmembrane helix of the mammalian Na+/H+ exchanger important for drug recognition and cation translocation. J. Biol. Chem. 276, 43792–43800 (2001).

33. Slepkov, E.R., Rainey, J.K., Sykes, B.D. & Fliegel, L. Structural and functional analysis of the Na+/H+ exchanger. Biochem J. 401, 623–633 (2007).

34. Nakanishi, H. et al. Transport cycle of plasma membrane flippase ATP11C by cryo-EM. Cell Rep. 32, 108208 (2020).

35. Zhuang, Y. et al. Structural insights into the human D1 and D2 dopamine receptor signaling complexes. Cell 184, 931–942 (2021).

36. Wang, L., Wu, D., Robinson, C.V., Wu, H. & Fu, T.M. Structures of a complete human V-ATPase reveal mechanisms of its assembly. Mol. Cell 80, 501–511 (2020).

37. Xiao, P. et al. Ligand recognition and allosteric regulation of DRD1-Gs signaling complexes. Cell 184, 943–956 (2021).

38. Hao, J. et al. Synthesis and pharmacological characterization of 2-(2,6-dichlorophenyl)-1-((1S,3R)-5-(3-hydroxy-3-methylbutyl)-3-(hydroxymethyl)-1-methyl-3,4-dihydroisoquinolin-2(1H)-yl)ethan-1-one (LY3154207), a potent, subtype selective, and orally available positive allosteric modulator of the human dopamine D1 receptor. J. Med. Chem. 62, 8711–8732 (2019).

39. Li, J. et al. Cryo-EM structures of *Escherichia coli* cytochrome *bo3* reveal bound phospholipids and ubiquinone-8 in a dynamic substrate binding site. Proc Natl Acad Sci USA 118, e2106750118 (2021).

40. Toelzer, C. et al. Free fatty acid binding pocket in the locked structure of SARS-CoV-2 spike protein. Science 370, 725–730 (2020).

41. Andreini, C., Bertini, I., Cavallaro, G., Holliday G.L. & Thornton, J.M. Metal ions in biological catalysis: from enzyme databases to general principles. J Biol Inorg Chem 13, 1205–1218 (2008).

42. Frenz, B., Walls, A., Egelman, E., Veesler, D. & DiMaio, F. RosettaES: a sampling strategy enabling automated interpretation of difficult cryo-EM maps. Nat Methods 14, 797–800 (2017).

43. Frenz, B. et al. Automatically fixing errors in glycoprotein structures with Rosetta. Structure 27, 134–139 (2019).

44. Moriarty, N.W., Grosse-Kunstleve, R.W. & Adams, P.D. electronic Ligand Builder and Optimization Workbench (eLBOW): a tool for ligand coordinate and restraint generation. Acta Cryst. D65, 1074–1080 (2009).

45. Wang, J., Wang, W., Kollman P. A. & Case, D. A. Automatic atom type and bond type perception in molecular mechanical calculations. Journal of Molecular Graphics and Modelling 25, 247260 (2006).

46. Jakalian, A., Bush, B.L., Jack, B.D. & Bayly, C.I. Fast, efficient generation of high-quality atomic charges. AM1-BCC Model: I. Method. J. Comp. Chem. 21, 132–146 (2000).

47. Greer, J. Three-dimensional pattern recognition: An approach to automated interpretation of electron density maps of proteins. J. Mol. Bio. 82, 279–301 (1974).

48. Pettersen, E.F. et al. UCSF Chimera--a visualization system for exploratory research and analysis. J. Comput. Chem. 25, 1605–12 (2004).

49. Wickham, H. ggplot2: Elegant graphics for data analysis. Springer-Verlag New York (2016). https://ggplot2.tidyverse.org

