## Supplemental Figures for "Automatic and accurate ligand structure determination guided by cryo-electron microscopy maps"

### ONLINE METHODS

#### *Creating protein-ligand dataset*

All single-particle EMDB entries with an associated ligand bound structure at 6Å resolution or better as of September 03, 2021 were obtained. Given the specificity of trying to model ions and glycans, structures with only these types of ligands were excluded from the dataset. Additionally, the set had several cases with small molecules in close proximity. To simplify the docking situation, entries with 2 or more ligands within the binding pocket as defined in our docking protocol were also eliminated from the set. To only have entries that fit the EM density well, structures with a density correlation below 0.4 or that left large regions of density unmodeled were dropped. When considering the first instance of a unique ligand for each EMDB entry, there were a total of 1704 total cases to process for docking.

#### *File preparation for docking*

For accurate ligand docking, small molecules need proper protonation states and partial charges. SDF files of the first instance of each unique ligand-entry pair were downloaded from the PDB and hydrogen atoms were added using phenix.elbow [44]. With the protonation state assigned, a mol2 file with AM1-BCC partial charges was generated with antechamber [45, 46]. Finally, a Rosetta specific parameters file was created for each ligand. Receptors were cleaned by eliminating non-macromolecular atoms in the PDB file and replacing modified residues with their unmodified correspondent. The ligand to be docked was added to its position in the deposited structure and randomly translated 0-2Å in any direction before docking.

#### *Density erosion and alignment*

To ensure the quality of ligand conformations in the ligand pool during the genetic algorithm, randomly perturbed ligands were aligned into unmodeled density to generate the initial pool. Voxels in the density map within 10Å of the center of mass of the ligand but greater than 2.5Å from an atom in the receptor were searched and eroded in a modified erosion algorithm from previously described methods [13,47]. Blobs of density are often discontinuous and difficult to separate from noise at lower resolution. To account for the low resolution, skeletonizing density was performed in two successive steps with increasing strictness on erosion. On the first pass, peaks in the density were detected and eroded only considering voxels sharing a face with each other. This keeps connections between density blobs that may be disjointed. The remaining voxels were clustered into potential skeleton networks by separating groups of voxels that are 3Å away from another group. Only the largest network of voxels was chosen for further erosion to eliminate noisy voxels. The largest group of voxels underwent a second, stricter erosion that considered all voxels that share a face or edge with each other, leading to a pseudo-atomic skeleton.

The skeleton was used during initial ligand conformer generation of the genetic algorithm to ensure a starting pool of ligands that already fit into the density. Small molecules were randomly translated and perturbed in the binding pocket and half of the small molecules in the initial pool were "aligned" to the skeleton. For alignment, the ligands were centered on the center of mass of the skeleton, and then atom-skeleton point pairs were determined. The shortest distance of an atom-skeleton pair while searching over all pairs was found, and this search was repeated until either all atoms or all skeleton points had a unique pairing. For the coordinates in each atom-skeleton pair, a "topped out" harmonic function was used to restrain ligand atoms:

$$E_{ij} = 36(1 - e^{-{x_{ij}}^2/9})$$

where E is an energy penalty applied and x is the distance in Angstroms between the atom-skeleton pair. The ligand is aligned into the density over 2 stages of energy minimization with 20 and 15 short rounds of minimizations with the atom-skeleton restraints updated after each round.

##### *Docking protocol and analysis*

An initial population of 100 ligands were generated by randomly perturbing across a six-fold axis and the torsion angles of the ligand to be docked. Only half of the initial ligands were aligned to the density as described above to ensure diversity in the initial population. The ligand population was optimized over 10 generations of a genetic algorithm using default parameters in GALigandDock and a scoring function with a high electron density score weight of 100 to evaluate a ligand's fit into density. The top 20 ligand conformers at the end of the GA were further optimized using a cartesian minimization in Rosetta. Example scripts for running density-guided ligand docking are provided below.

All entries were run in triplicate and the lowest-energy model for each individual run was further analyzed for docking success. Only cases with 25 or fewer torsion angles were analyzed as the search space of ligands with more torsions becomes difficult to fully explore during a GA. This, along with losing cases from inherent failure during ligand processing, left 1053 cases to analyze. Because of a low confidence in the reference models due to their low resolution, docked models were not directly compared to their respective reference models. Instead, all reference models were relaxed into their EM density map in Rosetta using the cartesian minimization used after the genetic algorithm. Along with a symmetry-independent RMSD value, docked models were compared to reference models by the number of residues that make hydrogen bonds with the ligand and a density correlation calculated in Rosetta. These metrics were used to categorize docking results as matches (docked pose within 1Å of relaxed reference model); non-match, similar quality (>1Å RMSD, density $\text{correlation}_{\text{dock}}$ - density $\text{correlation}_{\text{deposited}} > 0.025$ and hydrogen bonds$_{dock}$ - hydrogen bonds$_{deposited}$ > -1); or non-match, worse quality (>1Å RMSD, density correlation$_{dock}$ - density correlation$_{deposited}$ < -0.025 or hydrogen bonds$_{dock}$ - hydrogen bonds$_{deposited}$ < -1). Further support for docking success was calculated by determining the convergence of lowest energy ligand models across the triplicate runs. The distance between atom pairs across models were calculated and results were further divided into those with 2 or more trajectories having their lowest energy models within 1Å RMSD, more than within 1Å for 60% of atoms, or within 1Å for fewer than 60% of atoms.

The following command in Rosetta was used for the low-pass filter of map EMDB 30475:

```
$ROSETTA/main/source/bin/density_tools.default.linuxgccrelease -truncate_hires 4.0 -
mapfile emd_30475.map -truncate_map
```

##### *Comparison of docked and EM models to crystal structures*

For each ligand-protein pair in the EMDB dataset, the PDB was searched for structures solved by X-ray crystallography at 2.6Å resolution or better with at least 50% sequence identity to the protein and containing the same ligand. Results from the PDB were filtered further to only contain entries with similar ligand binding pockets as the corresponding EM model. The crystal models were aligned to the EM models by aligning all residues within 10Å of the ligand using matchmaker in UCSF Chimera [47]. Once aligned, the density correlations of the ligands in the crystal models were calculated in Rosetta. All entries with a pocket-aligned RMSD greater than 1.5Å and a ligand density correlation lower than 0.1 of the EM model were discarded for being too unalike. This gave 129 ligand-bound EMDB structures with similar crystal models. The 100 cases from this set where EMERALD converged on the same model were analyzed by RMSD to the ligand in the aligned crystal model.

##### *Visual analysis and images*

Figures of ligand-bound models and their EM maps were created using UCSF Chimera [48]. Maps displayed in figures were changed to 1.0Å using the vop command in Chimera for visual consistency. Plotting of data was performed using the ggplot2 package in R [49].

##### *Example xml for docking:*

```
<ROSETTASCRIPTS>
  <SCOREFXNS>
          <ScoreFunction name="relaxscore" weights="beta_genpot">
                  <Reweight scoretype="elec_dens_fast" weight="100"/>

                  <Reweight scoretype="gen_bonded" weight="1.0"/>
                  <Reweight scoretype="coordinate_constraint" weight="1.0"/>
        </ScoreFunction>
  </SCOREFXNS>

  <MOVERS>
          <SetupForDensityScoring name="setupdens" />
          <LoadDensityMap name="loaddens" mapfile="%%map%%" />
```

```
        <GALigandDock name="dock" scorefxn="relaxscore" ngen="10" npool="100"
rmsdthreshold="1.0" smoothing="0.0" ramp_schedule="0.1,1.0" grid_step="0.325"
padding="5.0" nativepdb="%%native%%" sidechains="auto" final_exact_minimize="bbsc"
random_oversample="100" use_pharmacophore="false" skeleton_threshold_const="5.0"
neighborhood_size="7" sample_ring_conformers="1" reference_pool="map"/>
  </MOVERS>
  <PROTOCOLS>
        <Add mover="setupdens"/>
        <Add mover="loaddens"/>
        <Add mover="dock"/>
  </PROTOCOLS>
  <OUTPUT scorefxn="relaxscore"/>
</ROSETTASCRIPTS>
```

*Example command line for docking*

```
$ROSETTA/main/source/bin/rosetta_scripts.linuxgccrelease \
    -in:file:extra_res_fa $ligand_params_file \
    -in:file:overwrite_database_params \
    -gen_potential \
    -database $ROSETTA/main/database \
    -score::gen_bonded_params_file
scoring/score_functions/generic_potential/generic_bonded.round6p.txt \
    -s $input_model \
    -overwrite \
    -multi_cool_annealer 10 \
    -parser:protocol $xml_file \
    -parser:script_vars map=$em_map \
    -atom_mask 2 \
    -sliding_window 1 \
    -edensity::score_sliding_window_context \
    -edensity::mapreso $reso \
    -edensity::grid_spacing 1.0 \
    -no_autogen_cart_improper
```

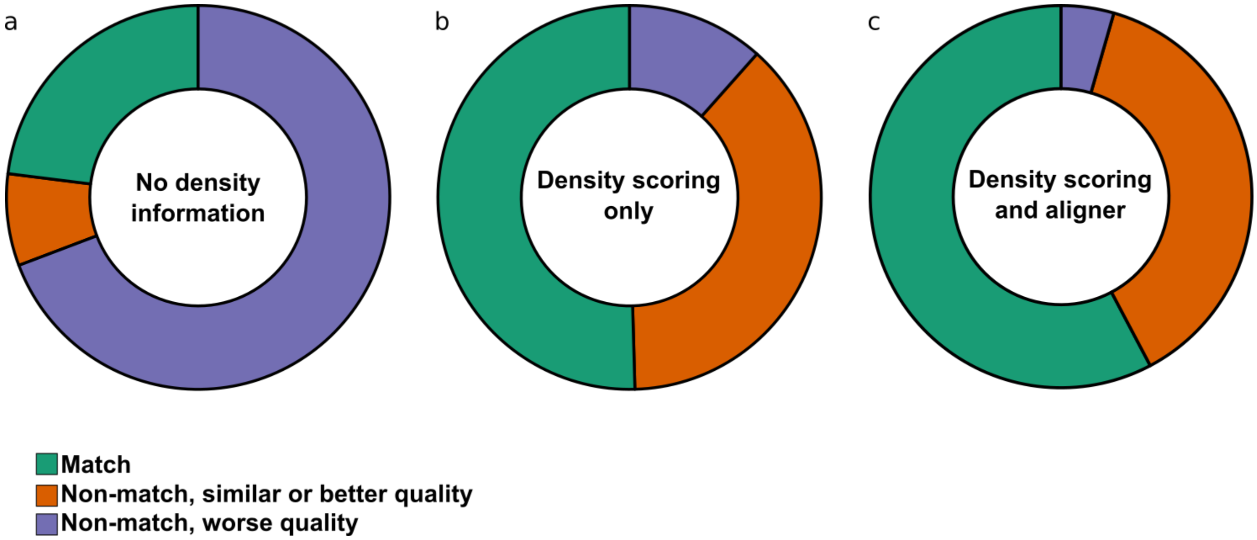


**Supplemental Fig. 1. Docking results with varying density information.** Docking results when docked with no density information in the genetic algorithm (a) and with density scoring evaluation during the genetic algorithm, but not during sampling (b). (a) Without density information, ligand docking can produce a model within 1Å RMSD to the deposited model for only 23% of cases, and models often do not fit the density, having a worse density correlation or fewer hydrogen bonds for 69% of cases. (b) Adding a density fit score during GA evaluation greatly improves modeling success, getting 50% of cases within 1Å RMSD to the deposited structure, but is still lower than when density information is included in sampling (c).

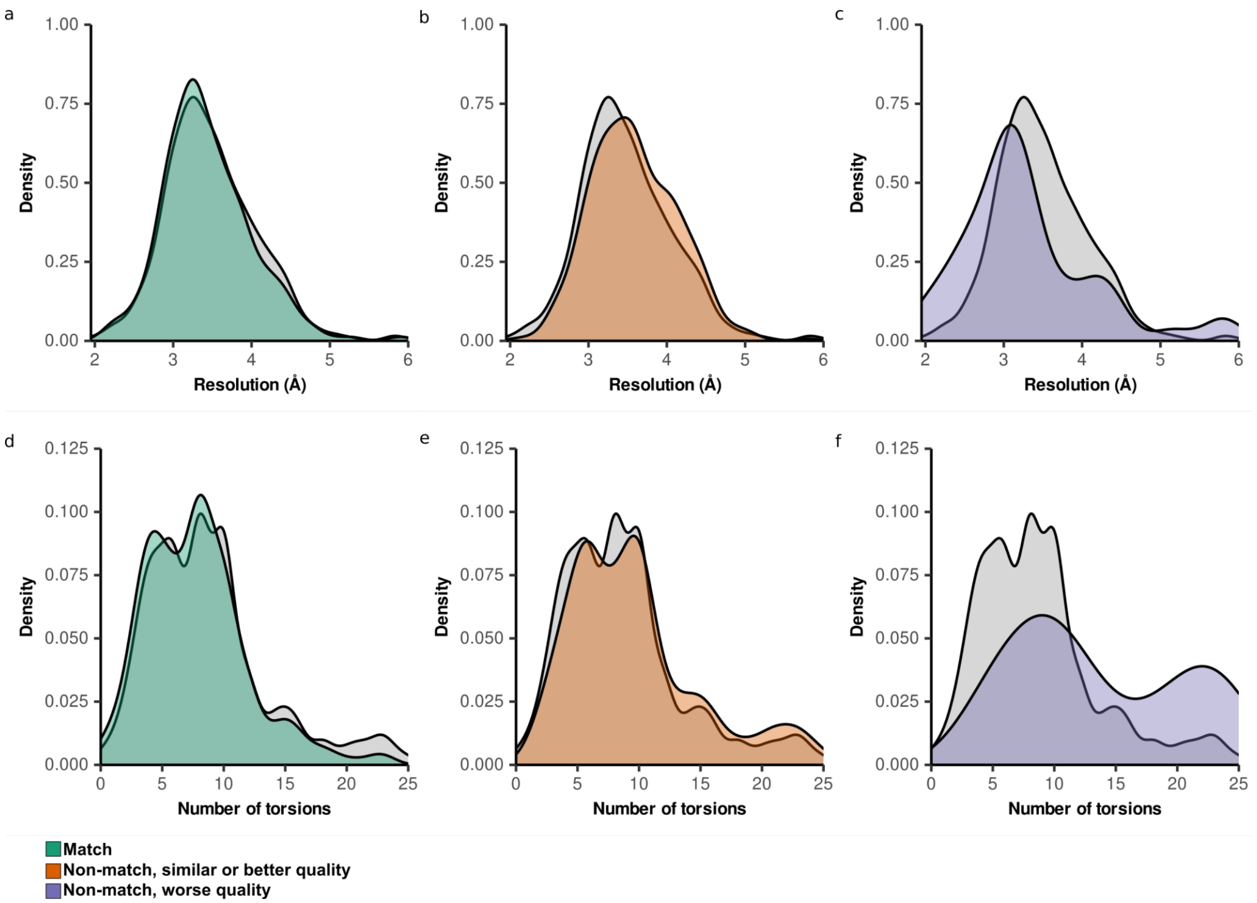


**Supplemental Fig. 2. Distribution of entry features by docking result.** (a-c) Resolution density plots for each docking categorization (*colors*) compared to the distribution of the entire dataset (*gray*). EM map resolution distributions for matched and docked models with similar quality do not vary from the whole dataset (a, b). Docked models with worse quality are skewed towards higher resolution maps, but this is likely because of variations in local resolution around the ligand or higher resolution maps being more likely to model more flexible ligands like lipids. (d-f) Density plots of the number of torsions in the docked ligand for each docking categorization (*colors*) compared to the distribution of the entire dataset (*gray*). The matched and similar quality docked models follow the distribution of the entire dataset (d, e). Cases where we produce a worse model disproportionally have ligands with a high number of torsion angles (f).

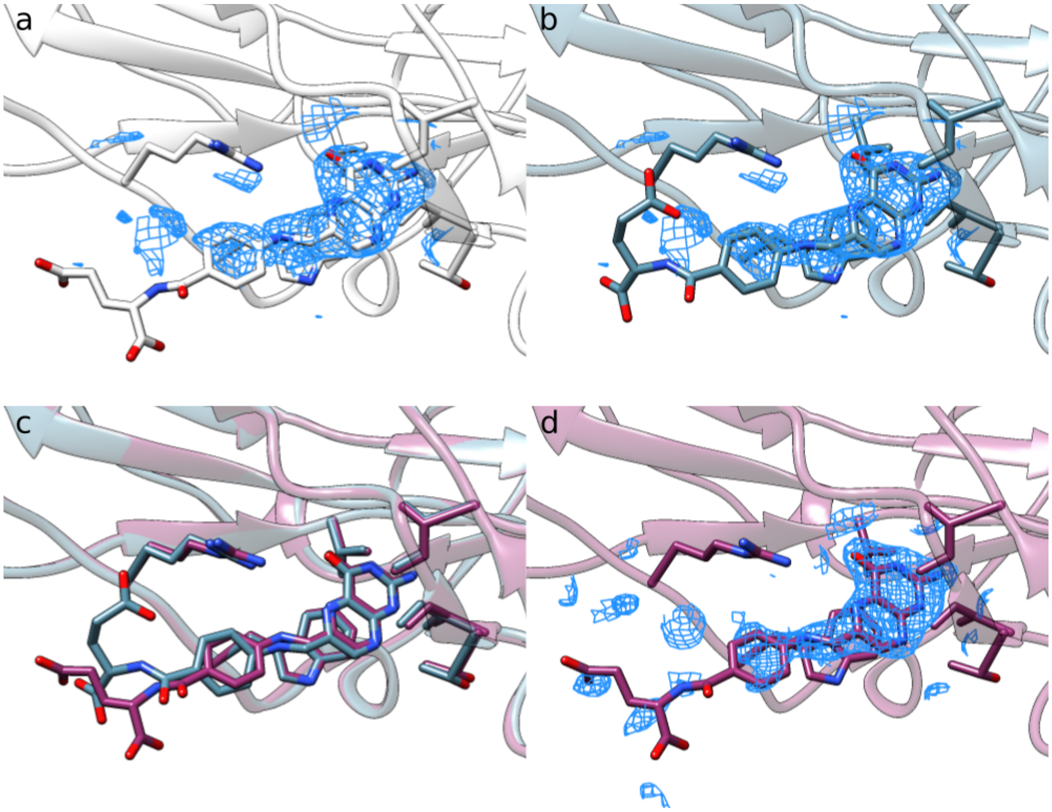


**Supplemental Figure 3. Ligand models of folate in MERS-CoV.** (a,b) The deposited model (EMDB: 23674, PDB: 7M5E) (a) and docked model (b) are similar in ordered regions of the ligand with strong cryoEM density and vary in the solvent exposed unordered region. (c) A superposition of the docked model (*blue*) and crystal model (*purple*) reveals alignment where the ligand is interacting with the receptor but disagreement in the unordered region. (d) The crystal model shown with its 2mFo-Fc map (contour = 1σ) shows weak density in the solvent exposed region.

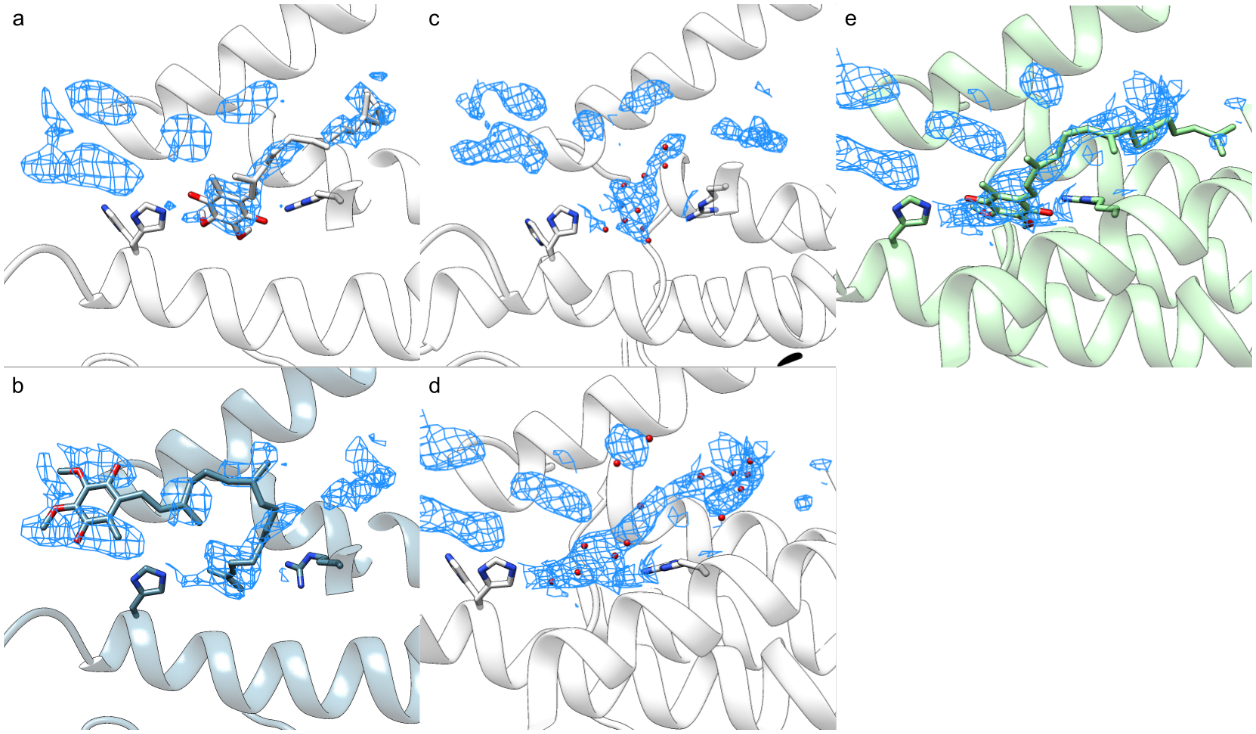


**Supplemental Fig. 4. Low-pass filtered map leads to a more complete skeleton and produces a model similar to the deposited model.** (a,b) Ubiquinone binding site in cytochrome bo3 in the deposited model (a) and docked model (b). EMERALD cannot find the known binding conformation and places the ligand in noisy density. (c) The density skeleton determined by our erosion protocol only includes density for a small portion of the ligand. (d) When the EM map is low-pass filtered at a 4Å cutoff, the density is more continuous, and the skeleton covers all of the ligand density. (e) The lowest-energy model from EMERALD after the map processing has a similar density correlation and similar hydrogen bonds as the deposited model.
